# Adeno-associated virus (AAV)-TBX18 does not generate biological pacemaker activity, unlike AAV-Hcn2

**DOI:** 10.1101/2025.09.11.672351

**Authors:** Jianan Wang, Mathilde R. Rivaud, Mischa Klerk, Arie R. Boender, Ruud N. Visser, Rinske Sparrius, Hee Young Lee, Karel van Duijvenboden, Yuting Yang, Emiel J.M. Kramer, Kyung Ho Park, Larry C. Park, Silke Schrödel, Christian Thirion, Eric Ehrke-Schulz, Anja Ehrhardt, Osne F. Kirzner, Klaus Neef, Hanno L. Tan, Arie O. Verkerk, Vincent M. Christoffels, Gerard J.J. Boink

**Author notes:** Address for correspondence: Gerard J.J. Boink, Amsterdam University Medical Centers, Meibergdreef 9, 1105AZ, Amsterdam, the Netherlands. Conflict-of-interest statement: J.W., A.R.B., Y.Y., E.J.M.K., O.F.K., K.N., and H.L.T. are employees of PacingCure BV. O.F.K., H.L.T,. and G.J.J.B. report ownership interest in PacingCure BV. S.S. and C.T. were employees of Revvity Gene Delivery GmbH, a wholly-owned subsidiary of PerkinElmer Inc. C.T. receives shares from the company. H.Y.L, K.H.P., and L.C.P. are employees of Naason Science Inc. L.C.P reports ownership interest in Naason Science Inc. J.W., R.V., V.M.C., and G.J.J.B. filed patent applications concerning application of uORF technologies. Other authors declare no conflicts of interest.

## Abstract

Gene therapy-based biological pacemakers have been proposed as an alternative to their hardware-based counterparts. In this context, short-term ectopic expression of the T-box transcription factor 18 (TBX18) in the ventricle reportedly generated potent short-term pacemaker function in various animal models. Here, we investigated the effect of adeno-associated virus (AAV)-mediated long-term expression of TBX18, and compared the outcome to that of the pacemaker ion channel Hcn2. Our findings revealed that CMV-driven ectopic TBX18 expression in mouse hearts led to severe cardiac fibrosis. At lower, non-fibrogenic levels, TBX18 maintained its transcriptional function but failed to induce pacemaker phenotypes. TBX18-expressing cells showed suppressed expression of key working myocardial genes, but the pacemaker gene program was not induced. Electrophysiological studies showed abnormal automaticity in TBX18-expressing cells, combined with prolonged repolarization and various current changes. However, no hyperpolarization-activated funny current was detected. In a complete AV-block rat model, AAV-mediated Hcn2 expression induced robust ectopic pacemaker activity in the presence of isoproterenol, whereas TBX18 expression neither generated such activities, nor augmented Hcn2-mediated pacing. In conclusion, at functional non-fibrogenic levels, TBX18 is neither sufficient nor necessary to induce pacemaker activity. In contrast, Hcn2 generates reliable pacing, making it a more viable candidate for biological pacemaker development.

## Introduction

Symptomatic bradycardia can pose a significant risk to life if left untreated. Electrical pacemakers have been used to successfully manage this condition, effectively restoring and maintaining normal cardiac rhythm. However, shortcomings inherent to these devices, such as limited battery life, complications associated with surgical procedures and inadequate autonomic modulation, underscore the necessity for more advanced therapeutic strategies (1, 2).

Gene therapy-based biological pacemakers are a promising alternative to electronic pacemakers (1, 3, 4). By modulating gene expression to mimic function of sinoatrial nodal (SAN) pacemaker cells, biological pacemakers aim to induce spontaneous diastolic depolarizations, capable of driving the heart at physiologically relevant rate. One approach is to manipulate ion channels (e.g. Hcn2 and Kir2.1) in order to increase inward current and/or decrease outward current during diastole (5–9). Transduced cardiomyocytes would then spontaneously depolarize and function as pacemaker cells. A potentially more comprehensive approach is based on transcription factor overexpression to reprogram cardiomyocytes into pacemaker cells that are functionally and phenotypically similar to SAN pacemaker cells (10, 11). These reprogramed cardiomyocytes are expected to emulate SAN cells not only in their electrophysiological properties, but also in their morphology and gene expression profiles.

T-box transcription factor (TBX) 3 and 18 are required for the development of the SAN and as such represent obvious targets for the biological pacemaker development (12, 13). Ectopic atrial TBX3 expression was shown to impose pacemaker phenotype using a transgenic expression system activated during early embryogenesis (14). However, tamoxifen-induced TBX3 expression in adult cardiomyocytes failed to do so but solely suppressed working myocardial phenotype (10). On the other hand, adenoviral (AdV) vector-mediated TBX18 overexpression was reported to converted ventricular chamber cardiomyocytes into pacemaker cells and generated biological pacing in both small and large animal models of heart block (11, 15, 16). Moreover, TBX18 overexpression in subsidiary atrial pacemaker tissue was shown to increase the spontaneous beating rates (17). However, the AdV vector used in these studies is highly immunogenic and, as a result, provides only transient transgene expression, impeding clinical translation (18, 19). In line with this immunological issue, pacing function waned during the follow-up period of several weeks. Alternatively, this transient pacemaker phenotype could also be due to insufficient reprogramming. Direct injection of TBX18 mRNA was recently explored as an alternative, but this approach was complicated by the need transforming growth factor B (TGFB) inhibition (as a co-factor to suppress fibrosis), and remained suboptimal in terms of pacemaker durability (20).

Adeno-associated viral (AAV) vectors are non-pathogenic, have low immunogenicity and can sustain long-term expression lasting for years (4, 21). Systemic delivery of AAV vector has been used in the field of cardiac gene therapy for over a decade, providing robust transgene expression in the whole heart (22–25). Recently, local delivery of AAV vectors has been demonstrated by several groups to achieve efficient transgene expression in large animal models (26, 27). AAV vectors have been approved by the US regulatory authorities for human application, further illustrating the strong potential for clinical translation (28). We hypothesized that ectopic AAV-mediated TBX18 expression would convert chamber cardiomyocytes into pacemaker cells and thereafter maintain a stable pacemaker phenotype. In order to test this hypothesis, it was critical to reduce TBX18 expression to non-cytotoxic levels. This allowed us to conduct further mechanistic studies on TBX18-expressing cardiomyocytes that were not possible in previous studies due to the transient expression systems employed (11, 15, 20). These mechanistic studies indicated that TBX18 expression *in vivo* did not induce a pacemaker phenotype, but induced abnormal automaticity. In the absence of local tissue inflammation (that has confounded previous studies utilizing AdV vectors), this induced phenotype was unable to generate any meaningful pacemaker activity in a complete AV-block rat model, while AAV-Hcn2 generated reliable pacing in the presence of isoproterenol.

## Results

### Adenoviral vectors support ectopic pacing in rats with complete AV-block

Previous studies showed that AdV-mediated TBX18 overexpression in ventricular cardiomyocytes (CM) induced the formation of ectopic pacemaker activity (11, 15, 16). To be able to recapitulate these findings, we used an established surgical AV-block rat model (29, 30). Electrical AV node ablation was performed and the successful generation of complete AV-block was confirmed by disassociation between P waves and QRS complexes on the surface cardiac electrocardiogram (ECG) (Figure 1A). We injected AdV vector expressing FLAG-tagged human TBX18 (AdV-TBX18) or GFP (AdV-GFP) as a control intramyocardially to the apex of the heart (Figure 1B). Saline injection was used as sham control. One week later, ECGs were measured before and after isoproterenol administration. Successful transgene delivery was confirmed by immunohistochemistry (Figure 1C). ECG analyses (Figure 1D) revealed that both AdV-TBX18 and AdV-GFP animals had significantly more profound ectopic pacing from the injection site and higher heart rate following isoproterenol administration when compared to the saline group (Figure 1, E and F). However, no significant difference was observed between AdV-TBX18 and AdV-GFP animals regarding the frequency of the ectopic pacing or the heart rate (Figure 1, E and F). Picrosirius red staining showed extensive cardiac fibrosis at the injection site of both AdV-TBX18 and AdV-GFP animals (Figure 1, G and H). These results suggested that the ectopic pacing observed was likely based on local tissue inflammation generated by the AdV vector used, rather than by TBX18 overexpression.

**Figure 1.**
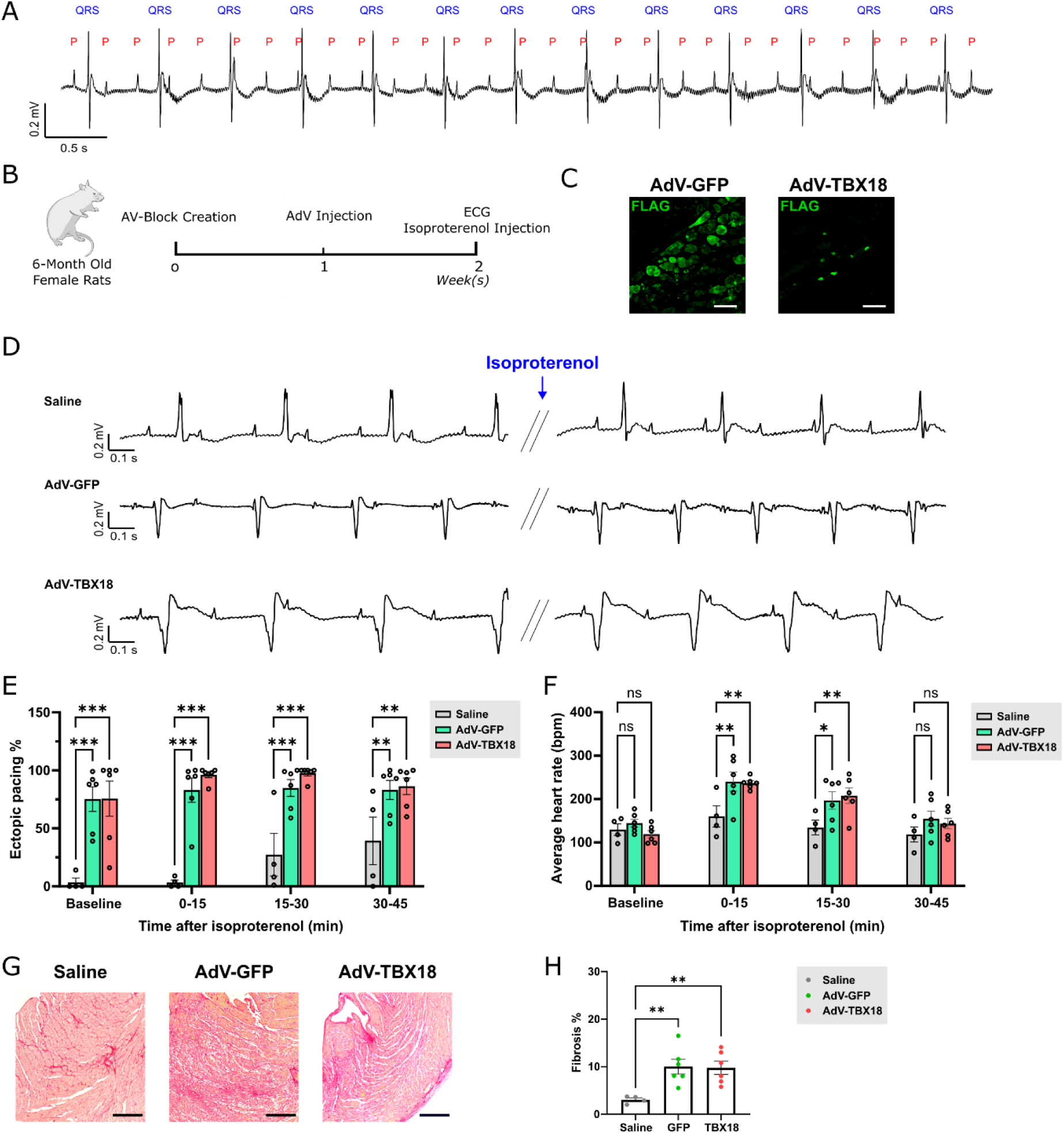
Injection of adenoviral vectors (AdV) leads to ectopic pacing in a complete atrioventricular block (AV-block) rat model. (**A**) An example ECG tracing showing complete AV-block in rats after AV node ablation. P wave and QRS complex are indicated on top. (**B**) Experimental design. (**C**) Immunofluorescence staining images of GFP and TBX18 in rat hearts injected with AdV-GFP and AdV-TBX18, respectively. Scale bar = 50 µm. (**D**) Example ECG tracings of rats injected with saline, AdV-GFP and AdV-TBX18, before and after the administration of isoproterenol. (**E**) Percentage of ectopic pacing and (**F**) average heart rate in rats injected with saline (n = 4), AdV-GFP (n = 6) and AdV-TBX18 (n = 6). (**G**) Picrosirius red staining images and (**H**) fibrosis quantification of rat hearts injected with saline (n = 4), AdV-GFP (n = 6) and AdV-TBX18 (n = 6). Scale bar = 300 µm. Data are shown as mean ± SEM. Data were compared using two-way ANOVA (E & F) or one way ANOVA (H) with *post-hoc* Fisher’s LSD test . *p < 0.05; **p < 0.01; ***p < 0.001; ns, not significant.

### TBX18 ectopic expression generates fibrotic scars in the mouse heart

To circumvent the use of AdV vector and at the same time achieve long-term TBX18 expression, we injected adult mice with AAV vectors containing CMV promoter and FLAG-tagged human TBX18 (AAV-CMV-TBX18). AAV vectors containing CMV promoter and FLAG-tagged GFP were used as control (AAV-CMV-GFP). To our surprise, a profound fibrotic scar was observed at the injection site in 9 out of 10 TBX18-injected mice, 4 weeks post injection, while such scar was absent from all 9 GFP-injected mice (Figure 2, A and B). Interestingly, the single TBX18 heart in which fibrosis was not detected was found to express TBX18 at the lowest level among the 10 TBX18 hearts (Supplemental Figure 1), suggesting that the fibrosis is likely related to the high levels of TBX18 overexpression, typically generated with conventional CMV-driven vectors. To investigate the progression of fibrosis, we performed histological studies on hearts harvested 4 days, 7 days and 28 days post injection of AAV-CMV-TBX18. Cardiac fibrosis appeared as early as 7 days post injection and progressed into substantial myocardial scars at 28 days post injection (Figure 2C). Immunofluorescence staining revealed that TBX18 expression accompanied the fibrotic tissues (Figure 2D). These results indicate that high levels of TBX18 overexpression are cytotoxic and lead to severe cardiac fibrosis.

**Figure 2.**
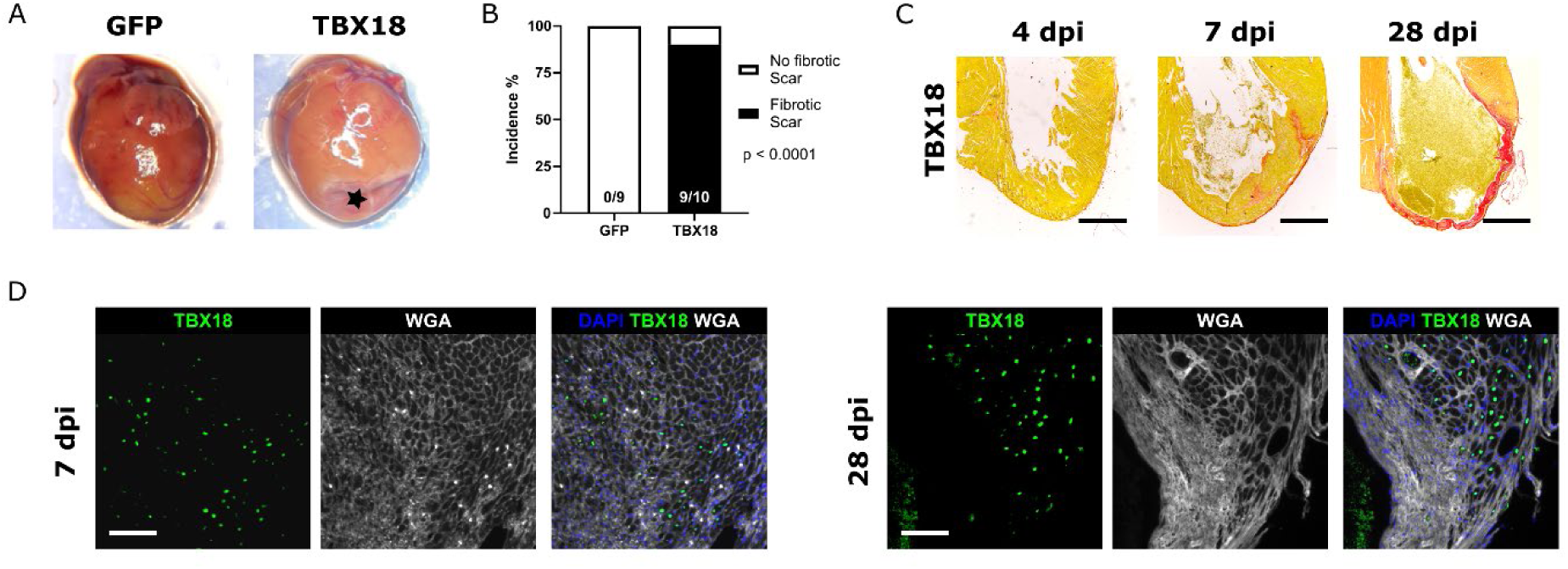
AAV-mediated TBX18 overexpression induces cardiac fibrosis in mouse hearts. (**A**) Macroscopy images of mouse hearts injected with AAV vectors overexpressing GFP or TBX18. Black star indicates the fibrotic scar at the injection site. (**B**) Incidence of hearts with fibrotic scar. Number of animals with fibrotic scar and total number of animals studied are presented within the associated bar charts. P value was calculated using Fisher’s exact test. (**C**) Picrosirius red staining images of mouse hearts injected with AAV-CMV-TBX18 at 4, 7 and 28 days post injection (dpi). Scale bar = 1 mm. (**D**) Immunofluorescence staining images of mouse hearts injected with AAV-CMV-TBX18 at 7 and 28 dpi. Scale bar = 100 µm.

### Optimized AAV vector expressing TBX18 circumvents fibrosis while maintaining transcriptional functionality

We hypothesize that the fibrosis observed was due to the supra-physiological levels of TBX18 expression. To reduce expression levels of TBX18, we added an upstream open reading frame (uORF) in the 5’ UTR of the transgene (Figure 3A), which was previously reported to reduce the translational efficiency.(31, 32) To test this approach in the context of an AAV expression cassette, we started with exploratory GFP (transfection) and TBX18 (transduction) experiments. As expected, introduction of this uORF did not change the transfection efficiency (fraction of GFP-positive cells), while effectively reducing GFP expression levels (Figure 3, B-D). Lowering the plasmid dose significantly reduced GFP expression level but also lowered the fraction of transduced cells (Figure 3, B-D). In a similar experiment, introduction of this uORF significantly reduced TBX18 expression level by 42% (Figure 3, E and F), confirming the effect of the uORF in our TBX18 overexpression vector. In neonatal rat ventricular myocytes (NRVM), introduction of the uORF and switching from the ubiquitously active CMV promoter to the cardiomyocyte-specific cTnT promoter reduced the TBX18 expression level to 25% and 39% to that of that from the original AAV-CMV-TBX18 vector, respectively (Figure 3, G and H). The combination of both further reduced the TBX18 expression level to 1% of the original vector and allows cardiomyocyte-specific expression (Figure 3G and 3H). We then sought to evaluate the function of this optimized AAV vector containing both cTnT promoter and uORF (AAV-cTnT-uORF-TBX18), which drives TBX18 at low expression levels and specifically in cardiomyocytes. NRVMs were transduced with both AAV-CMV-TBX18 and AAV-cTnT-uORF-TBX18 at a multiplicity of infection of 50,000. AAV-CMV-GFP was used as negative control while previously reported AdV vector bicistronically expressing TBX18 and GFP served as positive control (11). Quantitative real time-polymerase chain reaction (RT-qPCR) showed a significant upregulation of pacemaker gene *Hcn4* and down regulation of chamber genes *Gja1*, *Scn5a*, *Kcnj2*, and *Pln* in all three TBX18 vectors compared to the negative control. No significant differences were observed between cells transduced with AAV-CMV-TBX18 and AAV-cTnT-uORF-TBX18 (Figure 3I and 3J), indicating that the canonical transcription factor functionality of TBX18 was maintained in the optimized TBX18 vector.

**Figure 3.**
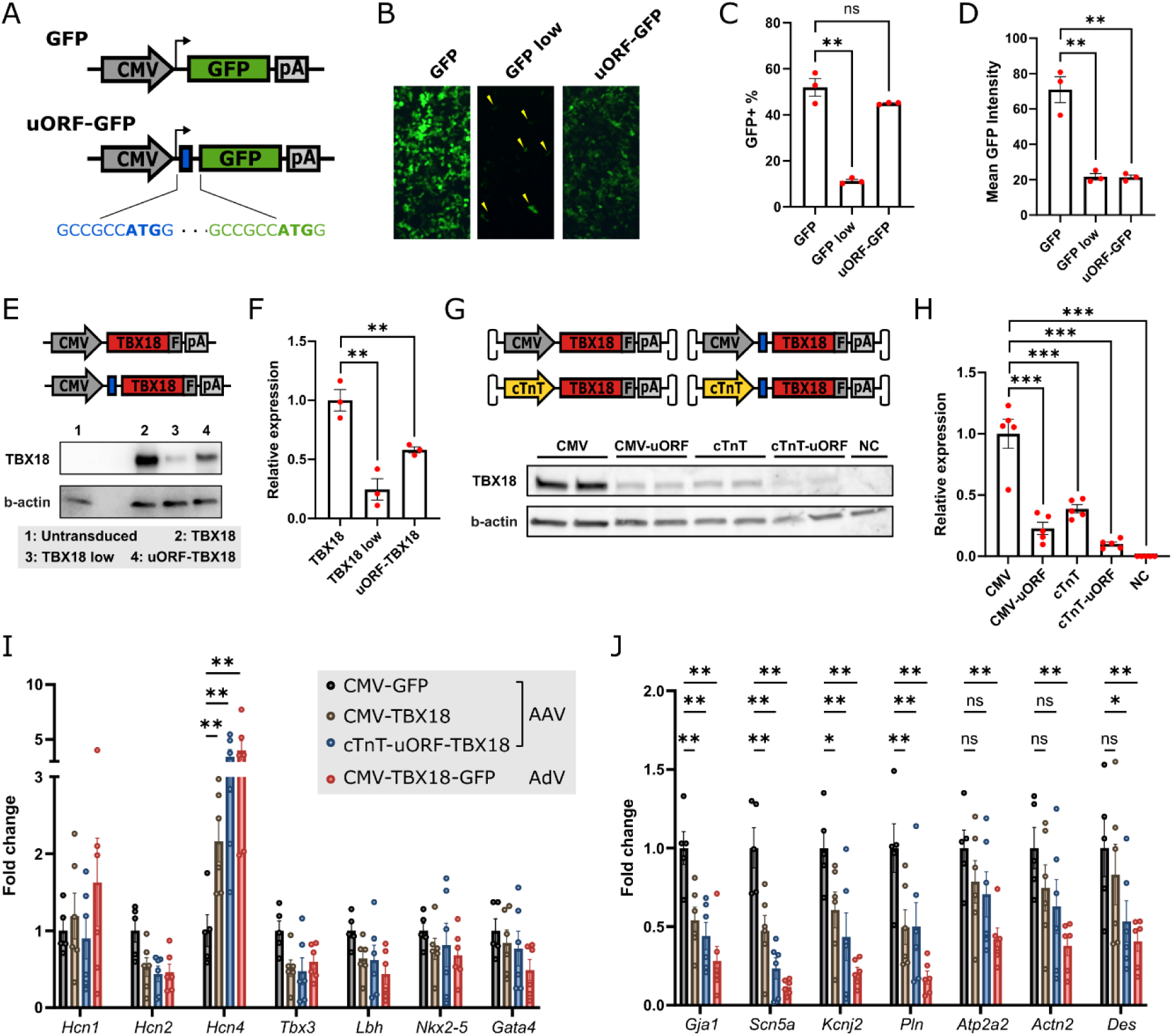
Dose-titrated expression cassette achieved low yet functional TBX18 expression. (**A**) Schematic representation of expression cassette with and without upstream open reading frame (uORF). Translation start sites are made bold. (**B**) Direct fluorescence images of HEK cells transfected with GFP and uORF-GFP plasmids. Yellow arrows indicate the GFP-positive cells. (**C**) Percentage of GFP-positive cells and (**D**) mean fluorescence intensity of HEK cells transfected with GFP or uORF-GFP plasmids (n = 3). (**E**) Example western blot and (**F**) quantification of TBX18 protein expression in HEK cells transfected with TBX18 or uORF-TBX18 plasmids (n = 3). (**G**) AAV vectors used for transduction and western blot of TBX18 protein expression in transduced cardiomyocytes. (**H**) Quantification of TBX18 protein expression in cardiomyocytes transduced with various TBX18 vectors (n = 3). (**I**) Expression of pacemaker genes and (**J**) working myocardial genes in cardiomyocytes transduced with various TBX18 vectors (n = 6) and one GFP vector (n = 5). Data are presented as mean ± SEM. Data were compared using Mann-Whitney test (H), one-way ANOVA with *post-hoc* Holm-Šídák test (C, D & F) or two-way ANOVA with *post-hoc* Holm-Šídák test (I & J). *p < 0.05; **p < 0.01; ns, not significant.

We subsequently studied fibrosis and myocardial scarring in mice injected with AAV-cTnT-uORF-TBX18, and compared outcomes to mice injected with the conventional AAV-CMV-TBX18 and two vectors expressing TBX18 incorporating either CMV-uORF or cTnT (without uORF; Figure 4A). Hearts injected with AAV-CMV-GFP were used as control. Picrosirius red staining illustrated that hearts injected with AAV-CMV-TBX18 formed an extensive fibrotic scar at the injection site with no viable myocardium left 4 weeks post injection (Figure 4B). On the contrary, hearts injected with AAV-cTnT-uORF-TBX18 had virtually no fibrosis, which was comparable to control (Figure 4, B and C). The other two groups showed intermediary outcomes. To study the long-term effect, we also performed analyses on hearts harvested 8 weeks post injection. Injections with AAV-CMV-TBX18 were not performed for this long-term study for animal welfare reasons. Fibrosis in hearts injected with AAV-cTnT-uORF-TBX18 did not differ from that in control hearts (Figure 4, D and E). These data indicated long-term tolerance of the AAV-cTnT-uORF-TBX18 vector. As AAV-cTnT-uORF-TBX18 vector is functionally equivalent to the other three vectors and tolerant for long-term expression, we continued our analyses with this vector.

**Figure 4.**
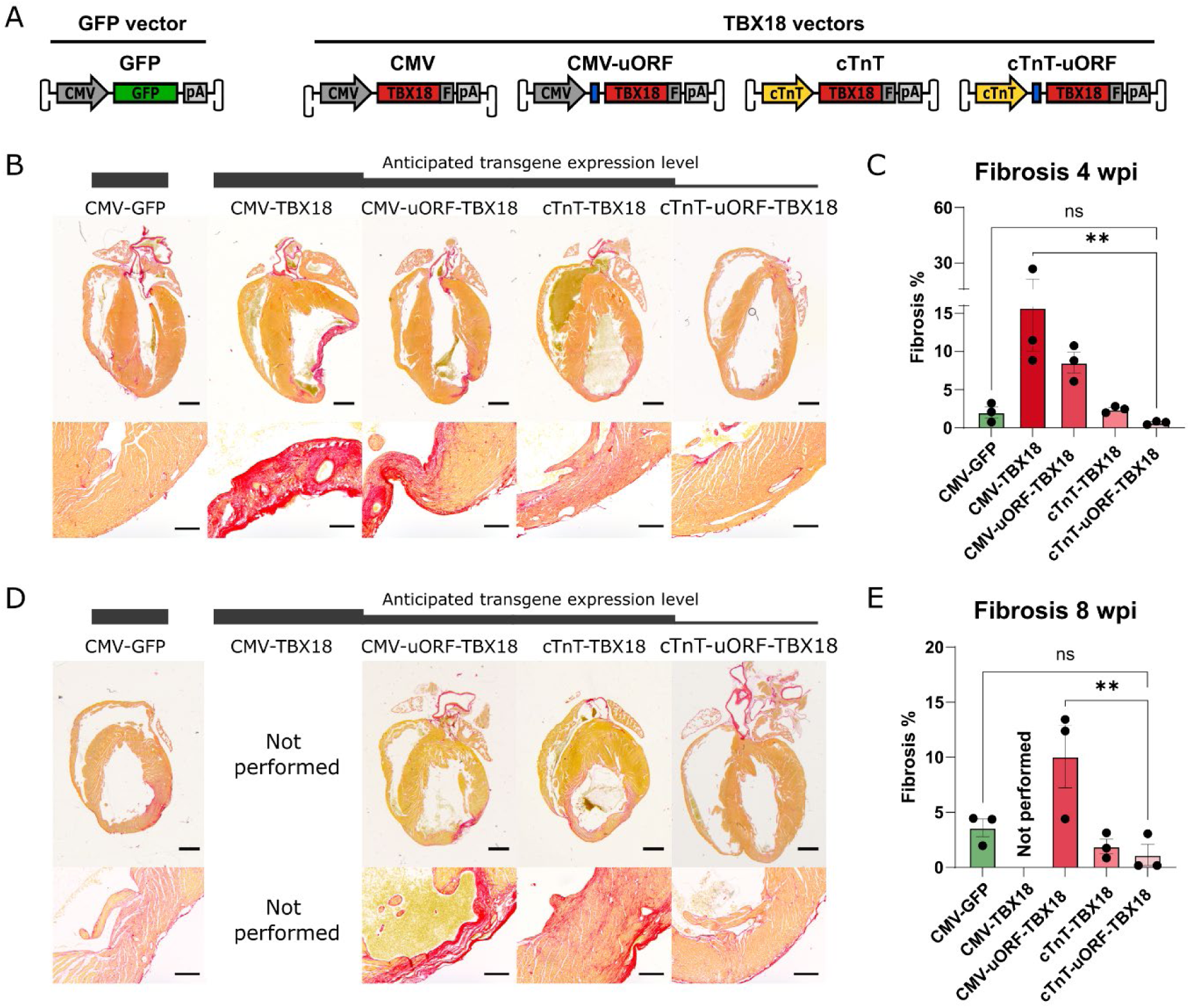
Long-term TBX18 expression delivered by cTnT-uORF-TBX18 vector does not induce cardiac fibrosis. (**A**) AAV vectors used for intramyocardial injection. (**B**) Picrosirius red staining images of GFP- or TBX18-injected mouse hearts 4 weeks post injection (wpi). Black bar at the top indicates the anticipated transgene expression level. Scale bar = 1 mm (upper) or 200 µm (lower). (**C**) Quantification of the cardiac fibrosis in the left ventricle of the injected hearts (n = 3). (**D**) Picrosirius red staining images of GFP-or TBX18-injected mouse hearts 8 wpi expressing GFP or TBX18. Black bar at the top indicates the anticipated transgene expression level. Scale bar = 1 mm (upper) or 200 µm (lower). (**E**) Quantification of cardiac fibrosis in the left ventricle of the injected hearts (n = 3). Data are presented as mean ± SEM. Data were compared using one-way ANOVA with *post-hoc* Fisher’s LSD test. **p < 0.01; ns, not significant.

### TBX18 suppresses the working myocardium phenotype

We focused the *in vivo* functional analysis on AAV-cTnT-uORF-TBX18, using AAV-CMV-GFP as a control (Figure 5A). Immunohistochemistry revealed expression of TBX18 or GFP at the injection sites, indicating robust transgene delivery (Figure 5, B and C). At 4 weeks post injection, Connexin43 (Cx43, encoded by *Gja1*), a direct downstream target of TBX18 (33) was efficiently suppressed in the TBX18-positive region when compared to the TBX18-negative region (Figure 5C). In control animals injected with AAV-GFP, Cx43 was homogenously detected across the GFP-positive and GFP-negative regions (Figure 5C). However, expression of the pacemaker ion channel Hcn4 remained undetectable at 4 or 8 weeks post injection, despite robust TBX18 expression (Figure 5B). As the positive control, Hcn4 was clearly detectable at the AV nodal region (Figure 5B).

**Figure 5.**
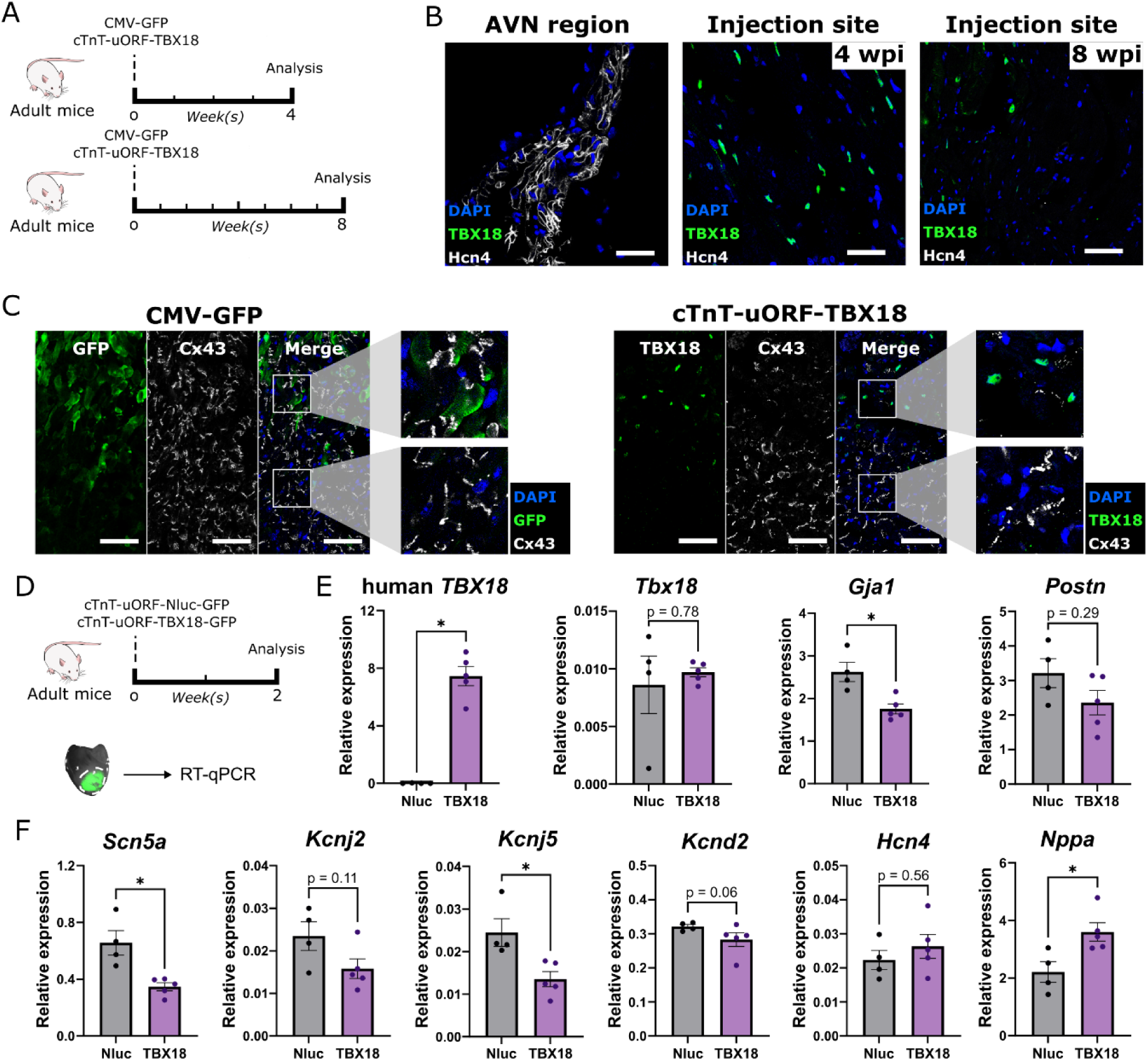
TBX18 expression at non-cytotoxic level suppresses working myocardial genes but does not induce pacemaker gene *Hcn4*. (**A**) Experimental design for histological analyses. (**B**) Immunofluorescence staining images of Hcn4 in TBX18-injected animals 4 and 8 weeks post injection (wpi). Scale bar = 50 µm. (**C**) Immunofluorescence staining image of Cx43 at injection sites of GFP- or TBX18-injected animals 4 wpi. Scale bar = 50 µm. (**D**) Experimental design for molecular analyses. (**E**) Expression level of human *TBX18*, *Tbx18*, *Gja1* and *Postn* determined by quantitative PCR (n = 4 for Nluc and n = 5 for TBX18). (**F**) Expression level of *Scn5a*, *Kcnj2*, *Kcnj5*, *Kcnd2*, *Hcn4* and *Nppa* determined by quantitative PCR (n = 4 for Nluc and n = 5 for TBX18). Data are shown as mean ± SEM. Data were compared using Mann-Whitney test. *p < 0.05.

To identify and dissect the transduced tissues, we modified AAV-cTnT-uORF-TBX18 by adding a self-cleaving p2A-GFP as a fluorescent marker (AAV-cTnT-uORF-TBX18-GFP) (Figure 5D). The same vector expressing nanoluciferase (AAV-cTnT-uORF-Nluc-GFP) was used as control. Two weeks post injection, the GFP-positive region was isolated by micro-dissection and used for quantitative RT-PCR to evaluate changes in gene expression. Expression of exogenous TBX18 was confirmed, whereas endogenous expression of *Tbx18* remained unaffected at low level (Figure 5E). A significant downregulation of the direct target *Gja1* indicates the transcriptional functionality of TBX18 delivered (Figure 5E). Expression of cardiac fibrosis marker *Postn* (34, 35) was not affected, confirming the non-cytotoxic expression level of TBX18 (Figure 5E). We observed significant down-regulation of the cardiac Na^+^ channel encoding gene *Scn5a* and K^+^ channel encoding gene *Kcnj5* (Figure 5F), which are important for cardiac conduction and modulation of resting membrane potential, respectively. K^+^ channel encoding genes *Kcnj2* and *Kcnd2* were non-significantly downregulated (p = 0.11 for *Kcnj2* and p = 0.06 for *Kcnd2*) (Figure 5F). On the other hand, *Hcn4* was not induced by TBX18 overexpression (Figure 5F), in line with our immunohistochemistry results. Ventricular stress marker *Nppa* was significantly upregulated (Figure 5F). RNA sequencing analysis confirmed these findings, and further strengthened the notion that working myocardial genes were suppressed but pacemaker genes were not induced (Supplemental Figure 2).

### TBX18 expression generates abnormal automaticity in mouse cardiomyocytes

To assess the electrophysiological consequences of TBX18 expression in cardiomyocytes, single left ventricular cardiomyocytes were isolated from mouse hearts 2 weeks post injection and were subjected to patch-clamp analysis. Spontaneous beating was clearly visible under the microscope in the isolated TBX18-expressing cardiomyocytes. Therefore, we proceeded to measure both spontaneous and overdrive-stimulated (4 Hz) action potentials (AP). Control cardiomyocytes did not generate spontaneous APs in 7 out of the 8 cardiomyocytes measured, while 8 out of the 8 TBX18-expressing cardiomyocytes exhibited spontaneous activity (Figure 6, A and B). The spontaneous APs of the TBX18-expressing cardiomyocytes had oscillations around plateau level and the time of such oscillations varied between approximately 2 to 10 seconds. We then performed 4 Hz overdrive stimulation to measure the AP parameters in detail. Typical recordings are shown in Figure 6C and average data are summarized in Figure 6D. All AP parameters measured differed between TBX18-overexpressing cardiomyocytes and control. The resting membrane potential of TBX18-expressing cardiomyocytes was depolarized by 5.8 mV, and the maximal AP upstroke velocity was significantly reduced, resulting in a significantly lower AP amplitude (Figure 6D). The AP durations at 20, 50 and 90% of repolarization were all significantly longer (Figure 6D).

**Figure 6.**
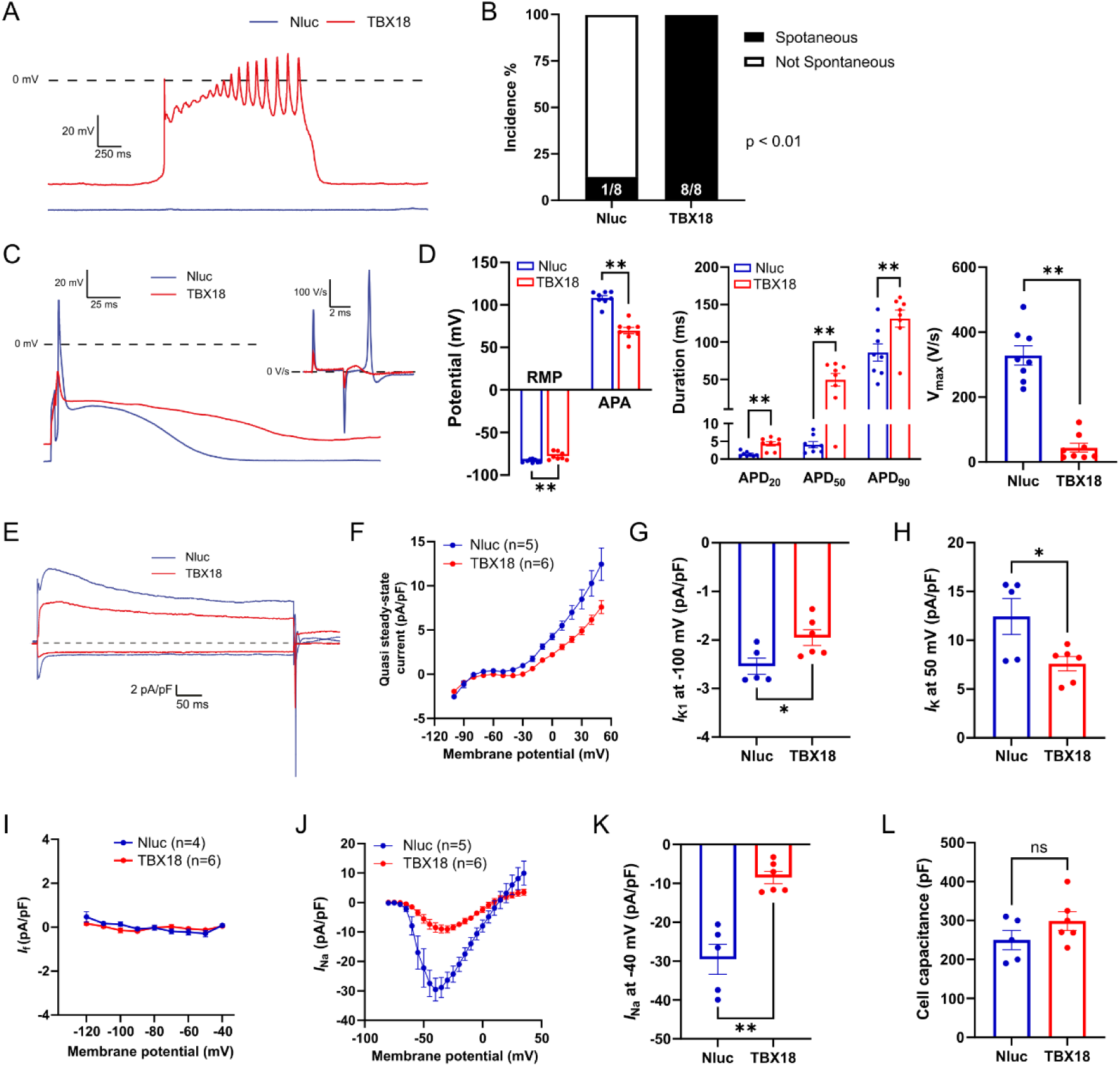
Long-term TBX18 expression at non-cytotoxic level leads to ionic remodeling of mouse cardiomyocytes. (**A**) Example action potentials (APs) recorded from TBX18-expressing and control cardiomyocytes (spontaneous / not stimulated). (**B**) Incidence of spontaneous activity. Number of cells with spontaneous activity and total number of cells studied are presented within the associated bar charts. (**C**) Example APs recorded from TBX18-expressing and control cardiomyocyte paced at 4 Hz. Inset: AP upstroke velocities. (**D**) Average AP parameters. RMP: resting membrane potential; APA: AP amplitude; APD_20/50/90_: AP durations at 20%/50%/90% repolarization; V_max_: maximal AP upstroke velocity (n = 8). (**E**) Example current traces at -100 and +50 mV of TBX18-expressing and control cardiomyocytes from a generic voltage clamp. (**F**) Current-voltage (I-V) relationships of the quasi steady-state current. (**G**) Current densities of the inwardly rectifying K^+^ current (*I*_K1_) measured at -100 mV. (**H**) Current densities of the delayed rectifier outward K^+^ current (*I*_K_) measured at 50 mV. (**I**) I-V relationships of the hyperpolarization-activated funny current (*I*_f_). (**J**) I-V relationships of Na^+^ current (*I*_Na_). (**K**) Peak *I*_Na_ density measured at -40 mV. (**L**) Cell capacitance reflecting cell size. (F-L) n = 4 for Nluc and n = 5 for TBX18. Data are shown as mean ± SEM. Data were compared using Fisher’s exact test (B) or Mann–Whitney test (D, G, H, K, L). *p < 0.05; **p < 0.01; ns, not significant.

We subsequently applied a generic voltage-clamp protocol to test net membrane currents in control and TBX18-expressing cardiomyocytes. Example current tracings and average voltage-current relationships indicate a difference in inward rectifier K_+_ current (*I*_K1_) and outward potassium currents (*I*_K_) between TBX18-expressing cells and control (Figure 6, E-H). Compared to control, TBX18-expressing cardiomyocytes presented a 23% reduction in *I*_K1_ (at -100 mV) (Figure 6G), coincident with the down-regulated *Kcnj2* expression found in our RT-qPCR analysis. TBX18-expressing cardiomyocytes also presented a 39% reduction in *I*_K_ (at 50 mV) (Figure 6H), coincident with the down-regulated *Kcnd2* expression found in our RT-qPCR analysis. In the voltage-clamp experiment, time-dependent, inward currents that activated upon the hyperpolarizing voltage steps were not observed, indicating absence of the funny current (*I*_f_) in both control and TBX18-overexpressing cardiomyocytes (Figure 6I). We next proceeded with Na^+^ current (*I*_Na_) measurement which revealed a 71.3% reduction in TBX18-overexpressing cardiomyocytes (Figure 6, J and K), consistent with the reduced maximal AP upstroke velocity and *Scn5a* expression. Whole-cell capacitance was also recorded as a measure of cell size, which was not different between TBX18-expressing and control cardiomyocytes (Figure 6L).

Taken together, our patch-clamp data revealed that ectopic TBX18 expression in mouse cardiomyocytes generates a distinctive electrophysiological phenotype that is incapable of sustaining a regular rhythm. The observed spontaneous depolarizations in these TBX18-expressing cardiomyocytes were the result of potassium channel dysregulation, rather than from the canonical mechanisms seen in SAN pacemaker cells.

### AAV-mediated Hcn2 expression generates ectopic ventricular pacemaker activity while TBX18 expression does not

Previous studies have suggested that interference with the inward rectifier current, via overexpression of a dominant negative variant of Kir2.1 can possibly liberate endogenous pacemaker activity (36), or support Hcn2-based pacemaker activity (8, 37). We therefore proceeded with testing AAV-cTnT-uORF-TBX18 in the complete AV-block rat model (Figure 7A). In doing so, we compared outcomes to overexpression of the pacemaker channel Hcn2 and also tested the combination of Hcn2 and TBX18 overexpression. Successful transgene delivery was confirmed via immunofluorescence staining (Figure 7B). At baseline, all animals showed similar frequency of ectopic pacing and average heart rate. Neither TBX18-injected animals nor saline-injected animals responded to the isoproterenol administration (Figure 7C-7E). However, intriguingly, Hcn2-injected animals exhibited robust ectopic pacing in response to isoproterenol (Figure 7, C and D), as illustrated by the increase in average heart rate (Figure 7E). Co-injection of AAV-cTnT-Hcn2 and AAV-cTnT-uORF-TBX18 did not further augment Hcn2-based pacemaker function (Figure 7, C-E). No significant cardiac fibrosis was observed at the injection site of both saline and AAV-injected animals (Figure 7, F and G). Taken together, our data shows that AAV-mediated Hcn2 expression generates a viable pacemaker phenotype in response to isoproterenol, while TBX18 expression does not.

**Figure 7.**
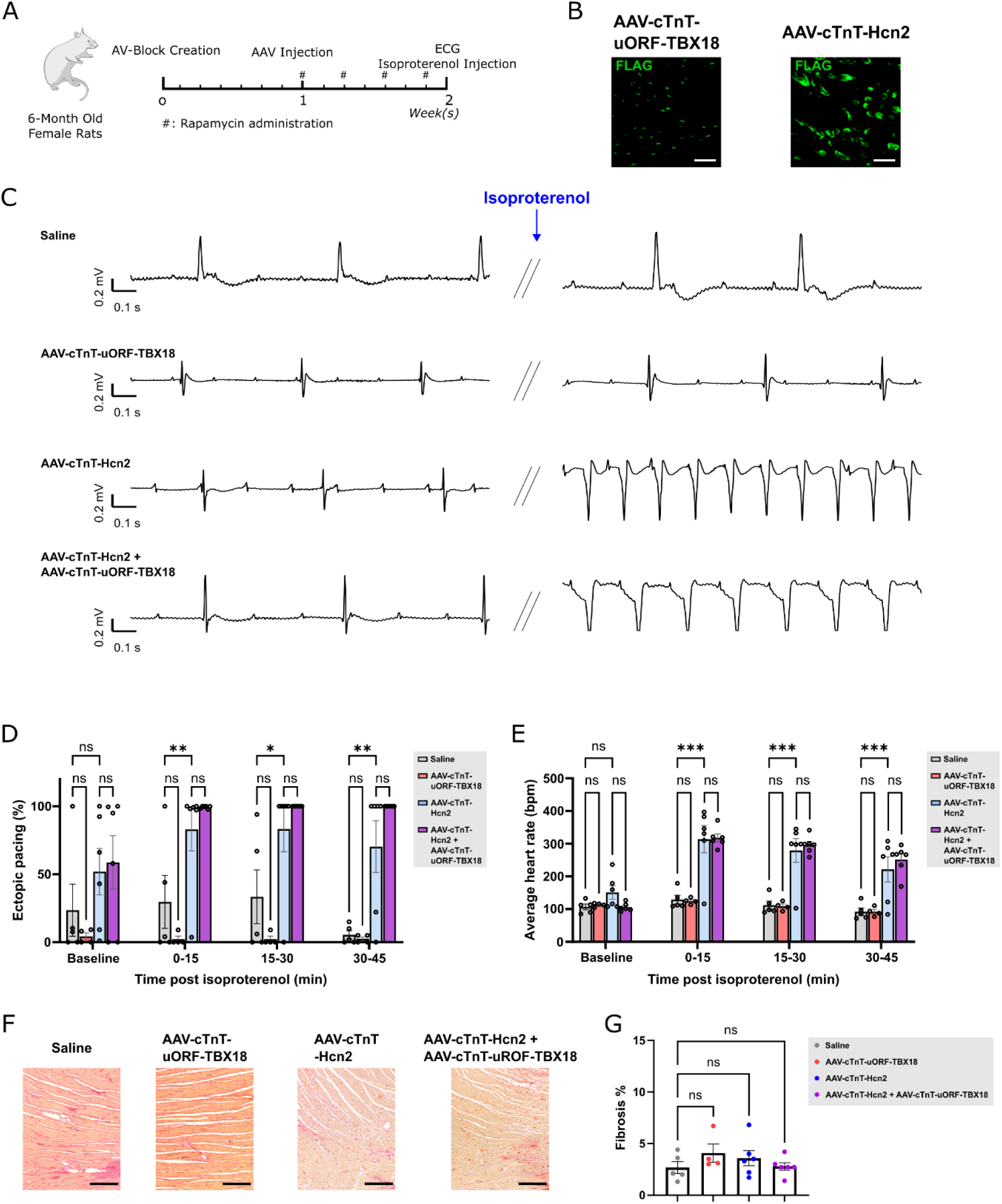
AAV-mediated expression of Hcn2 generates robust ectopic pacing in the presence of isoproterenol while AAV-mediated expression of TBX18 does not. (**A**) Experimental design. (**B**) Expression of TBX18 and Hcn2 in rat hearts injected with AAV vector expressing TBX18 and Hcn2, respectively. Scale bar = 50 µm. (**C**) Example ECG tracings of rats injected with saline or various AAV vectors before and after the administration of isoproterenol. (**D**) Percentage of ectopic pacing and (**E**) average heart rate in rats injected with saline or various AAV vectors . (**F**) Picrosirius red staining images and (**G**) fibrosis quantification of rat hearts injected with saline, AAV vector expressing TBX18, AAV vector expressing Hcn2, or both. (D-G) n = 5 for saline, n = 4 for TBX18, n = 6 for Hcn2 and TBX18 + Hcn2. Scale bar = 300 µm. Data are shown as mean ± SEM. Data were compared using two-way ANOVA (D & E) or one-way ANOVA (G) with *post-hoc* Fisher’s LSD test. *p < 0.05; **p < 0.01; ***p < 0.001; ns, not significant.

### Analysis the transcriptional changes induced by *Hcn2* and *TBX18* expression in NRVMs

To investigate the impact of *Hcn2* or *TBX18* expression on cardiomyocytes, we transduced NRVMs with the *Hcn2* vector and all four *TBX18* vectors used in this study, followed by RNA-sequencing. AAV vector with the cTnT promoter and no coding sequence was used as control. Principal component analysis revealed a clear segregation of the samples based on *Hcn2* and on *TBX18* expression. All samples of the same group clustered together, and all four TBX18 groups clustered closely based on overall expression profiles (Figure 8A). Differentially expressed genes across the groups clustered into three distinct expression profiles (Figure 8B). The Hcn2 and TBX18 groups shared similarity in gene expression profiles corresponding to gene cluster 1 and 2, where cluster 1 marks the upregulated genes and cluster 2 marks the down regulated genes compared to control. Notably, expression of the transcription factors necessary for SAN development (e.g. *Isl1*, *Shox2*, *Tbx3*) were not induced in any of the TBX18 groups (Figure 8, B and D), in line with our findings (Figure 3, Figure 5 and Figure 6) and other studies using NRVMs and mice (38, 39).

**Figure 8.**
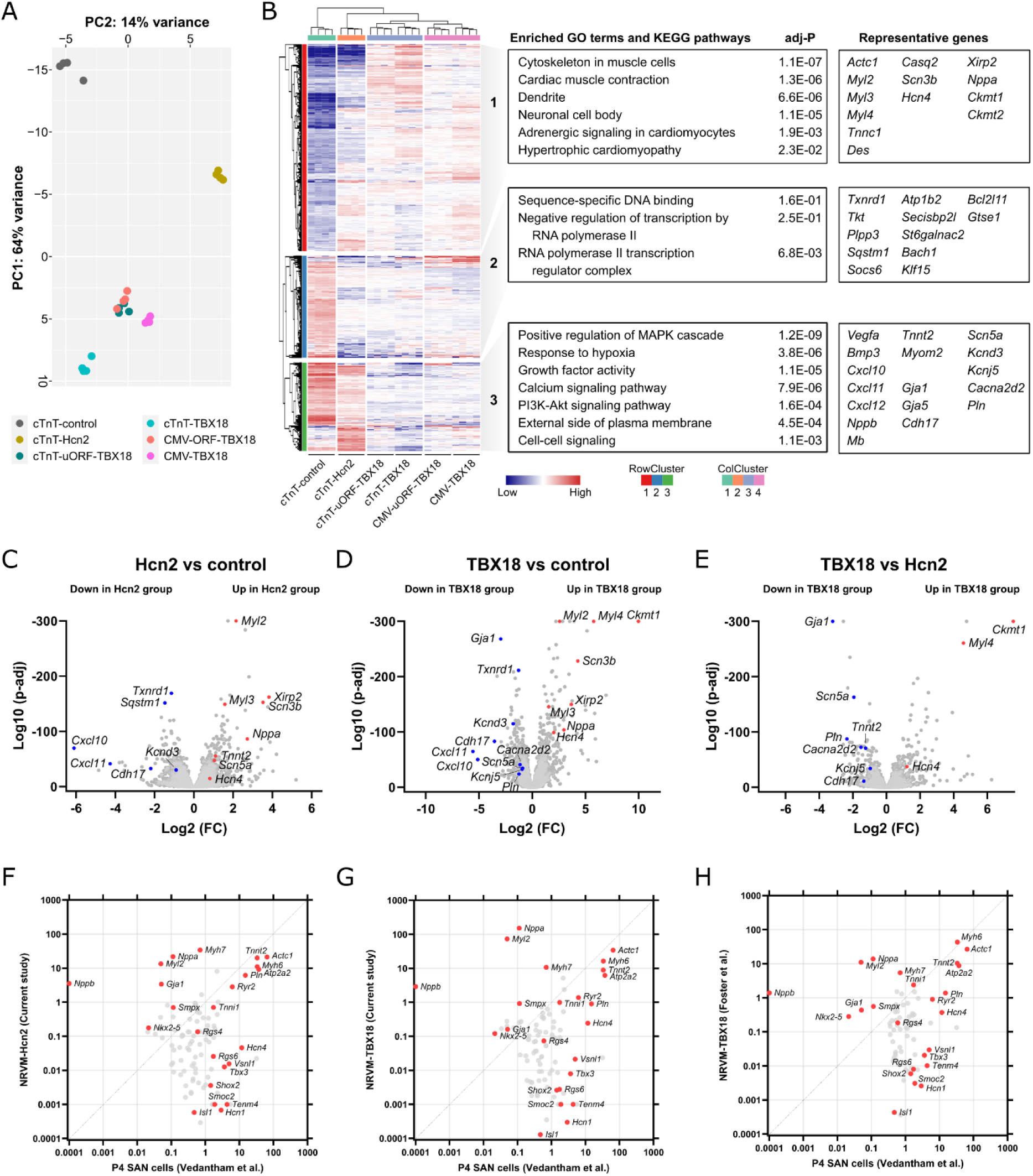
RNA sequencing analysis reveals the transcriptional effects of Hcn2 and TBX18 expression in NRVMs. (**A**) Principal component analysis of the cardiomyocytes transduced with control, Hcn2 and various TBX18 vectors. (**B**) Unsupervised functional annotation heatmap of NRVMs transduced with control, Hcn2 and various TBX18 vectors. (**C-E**) Volcano plots showing transcripts differentially expressed between NRVMs transduced with (C) control and Hcn2, (D) control and TBX18, and (E) Hcn2 and TBX18. (**F-H**) Scatter plots showing gene expression in NRVMs transduced with TBX18 or Hcn2 in comparison to postnatal day 4 (P4) mouse sinoatrial node (SAN) pacemaker cells. (F) Hcn2-transduced NRVMs from the current study (G) TBX18-transduced NRVMs from the current study. (H) TBX18-transduced NRVMs from Foster et al. (39).

Genes in cluster 3 were specifically suppressed in the TBX18 groups, and were related to various signaling pathways (e.g. *Vegfa*, *Bmp3*, *Pln*), ion channels (e.g. *Scn5a*, *Kcnj5*, *Cacna2d2*) and cell-cell communications (*Gja1*, *Gja5*, *Cadh17*) (Figure 8, B-E). Cluster 1 genes were induced in both TBX18 and Hcn2 groups compared to control, suggesting that the elevated expression of those genes may reflect a broader consequence of transgene overexpression, such as disruption of the normal maturation process and/or pathological remodeling of the ventricular myocytes. Notably, genes in this cluster, such as *Hcn4*, *Myl4* and *Nppa*, are typically expressed only in fetal, but not adult ventricular cardiomyocytes (40–42). We observed a 3-fold decrease in *Hcn4* expression over 5 days of culture in untreated NRVMs (Supplemental Figure 3), suggesting that the observed increase in *Hcn4* in TBX18 and Hcn2 groups compared to control may be caused by attenuated downregulation that normally occurs in untreated NRVMs during their maturation. Reactivation of these fetal genes in adult ventricular myocytes has been associated with cardiac disease and pathological remodeling (43–45).

We then compared the transcriptional profiles of NRVMs transduced with Hcn2 and TBX18 from the current study, as well as NRMV transduced with TBX18 from Foster et al. (39), to those of postnatal day 4 (P4) mouse SAN pacemaker cells (46) (Figure 8, F-H). Our analysis revealed that neither Hcn2 nor TBX18 induced a transcriptional signature resembling that of native SAN pacemaker cells. For example, the expression of the TBX18-induced level of *Hcn4* and *Isl1* remains 48-fold and 3600-fold lower than those in native newborn SAN pacemaker cells, respectively, whereas common cardiomyocyte genes were expressed at comparable levels. Comparison between these datasets to embryonic day 16.5 (E16.5) mouse SAN pacemaker cells (47) also showed similar results (Supplemental Figure 4). These results together suggest that transducing ventricular myocytes with Hcn2 or TBX18 does not result in the generation of *bona fide* SAN-like pacemaker cells. Furthermore, these data indicate the Hcn2-based biological pacing is likely attributable to the functional modification of working cardiomyocytes through overexpression of Hcn2 itself, as anticipated.

## Discussion

In this study, we ectopically expressed TBX18 in adult murine ventricular myocardium using AAV-mediated delivery with the goal to generate long-term pacemaker function (11, 15, 16). However, we observed that conventional overexpression of TBX18 leads to severe cardiac fibrosis, which could be prevented by modification of the expression cassette to reduce protein expression levels of TBX18 while maintaining its transcriptional function. We found that TBX18 expression at non-cytotoxic levels suppressed working myocardial genes, in accordance with previous findings (11, 15, 38), but did not induce a genuine pacemaker phenotype. As a result, AAV-mediated TBX18 expression did not induce a pacemaker phenotype in AV-block rats, in contrast to Hcn2 expression, which generated robust ventricular pacing in response to isoproterenol.

We observed ectopic pacing activities from both AdV-GFP-injected and AdV-TBX18-injected animals using a complete AV-block rat model, suggesting that these activities are induced by the AdV vector itself instead of the transgene delivered. Previous studies showed similar results where AdV-TBX18 generated automaticity in rat complete AV-block models however not significantly different from AdV-GFP-induced automaticity (29, 30). Moreover, AdV infection rapidly provokes inflammation and infiltration of host immune cells in hearts (48). Recombinant AdV vectors, albeit being replication-defective, still activate both innate and adaptive immunity, leading to an inflammatory response (18, 19, 49). Indeed, we and others found severe fibrosis at the injection sites of AdV-GFP and AdV-TBX18 (30). These immune/inflammatory responses make it difficult to specifically study the effect of candidate biological pacemaker genes, and might also contribute to the inconsistent outcomes in different studies utilizing AdV-TBX18 (11, 20, 30).

In our study, AAV vectors were used to deliver TBX18 because of their low immunogenicity, allowing us to study the effect of TBX18 overexpression without the confounding effects of a severe immune response. Moreover, the use of the AAV vector system also would enable long-term expression of TBX18, aiming to facilitate a chronic treatment effect. In our experiments, ectopic expression of TBX18 using a conventional CMV-driven AAV vector was highly cytotoxic and therefore fibrogenic in mouse hearts. The absence of fibrosis in control where GFP was expressed from the same vector ruled out other potential non-transgene related causes such as mechanical trauma from injection or immune responses to viral vectors. Indeed, others have also found that TBX18 induced significantly more fibrosis than GFP when expressed from directly injected mRNA or from AdV vectors (30, 50). Inhibition of TGFB signaling using small molecules has been reported to mitigate such fibrosis and maintains the TBX18-induced automaticity *in vitro* and *in vivo* (30). However, TGFB signaling is also heavily involved in epithelial to mesenchymal transition (51), which is the proposed mechanism of TBX18-mediated conversion from cardiomyocytes to pacemaker cells (39). Furthermore, Tgfb signaling plays a crucial role in many physiological processes, such as tissue development, homeostasis and injury repair (52). These pleiotropic effects of TGFB inhibition are therefore generally undesirable from a translational perspective.

In order to attenuate fibrosis, we lowered expression levels of TBX18. We did not lower the viral titer to reduce expression, because AAV transduction follows a Poisson distribution (53), hence lower transduction, will lead to a mosaic pattern of transduced and un-transduced cells, making it more difficult to reach the critical mass needed for reliable pacemaker performance. Instead, a uORF was inserted in the 5’UTR of the TBX18 expression cassette. The theoretical advantage of this approach is that it allows to reduce protein levels without the need to reduce transduction efficiency or transcriptional activity of the expression cassette. Both applying the uORF and switching to a cardiac-specific promoter that is less strong than CMV (54, 55) significantly reduced TBX18-induced fibrosis in mouse hearts, and the combination completely eliminated it. This combination reduced the expression level of TBX18 to approximately 1% of the original level but still maintained its transcriptional function, as the suppression of its direct downstream target *Gja1* (33) and other target genes remained robust both *in vitro* as well as *in vivo*. Our RNA sequencing results showed similar transcriptional profiles across all four TBX18 groups further supporting the functional equivalence of TBX18 delivered by different vectors.

Our *in vitro* and *in vivo* molecular experiments showed that ectopic expression of TBX18 suppressed the expression of chamber myocardial genes *Gja1*, *Scn5a, Kcnj5* and *Kcnj2*, in agreement with previous studies using a transgenic mouse line and AdV-mediated overexpression (11, 15, 38). This finding is further supported by our electrophysiological experiments where *I*_K1_, *I*_K_ and *I*_Na_ significantly decreased in TBX18-expressing cells. In terms of expression of pacemaker genes, neither the TBX18-transduced NRVMs from our study nor the ones from others (39, 56) show resemblance to the transcription profiles of the *bona fide* SAN pacemaker cells. For instance, genes for essential SAN pacemaker transcription factors ISL1, SHOX2 and TBX3, each essential for SAN pacemaker cell differentiation and maintenance, are not induced by TBX18 (57). Contradictory responses of *Hcn4* to ectopic TBX18 expression in NRVMs have been reported with two studies showing induction (11, 39) and another study showing reduction (56). In our study we observed higher expression of *Hcn4* in NRVMs transduced with TBX18 compared to control. However, a similar increase was also observed in NRVMs transduced with Hcn2, indicating the induction is not specific to TBX18 but may be a more general response to gene overexpression. The apparent induction of *Hcn4* expression levels may reflect the attenuated maturation of transduced NRVMs and failure to downregulate *Hcn4.* In fact, the induction of *Hcn4* by TBX18 has only been reported using neonatal cardiomyocytes (11, 20, 39) except one study using juvenile pig heart where improper reference tissue was used as control (15). In our adult mouse model we did not observed the induction of *Hcn4* expression at either mRNA or protein levels, nor did we find *I*_f_ in the TBX18-expressing cardiomyocytes. These results are consistent with those reported by Greulich *et al.,* in which misexpression of *Tbx18* in fetal mouse hearts suppressed the expression of working cardiomyocyte genes, but did not induce the expression of pacemaker genes (38). These results are also in line with the previous reports that TBX18 is primarily a transcriptional repressor rather than an activator (58, 59). Furthermore, the decrease in cell size was also considered as the result of the transition from mature cardiomyocytes to pacemaker cells (11), which we did not observe in the TBX18-expressing cardiomyocytes isolated from mice. Taken together, our results indicate that ectopic expression of TBX18 in cardiomyocytes is impacting on the expression of particular genes involved in ion handling that are expressed preferentially in working cardiomyocytes, thereby altering AP characteristics, but TBX18 is not converting NRVMs or adult rodent (i.e. mouse and rat) ventricular cardiomyocytes into pacemaker cells.

Interestingly, abnormal automaticity was observed in isolated TBX18-expressing cardiomyocytes, despite the lack of *I*_f_. These cells exhibited spontaneous depolarization, resulting in APs exhibiting early afterdepolarization (EAD)-like oscillation near the plateau. Since no *I*_f_ was detected in these cells, we interpret this spontaneous depolarization likely to stem from the decrease of *I*_K1_, which destabilizes the resting membrane potential. Indeed, suppression of *I*_K1_ by itself is enough to generate automaticity in cardiomyocytes, when achieved by overexpression of a dominant negative mutant of *Kcnj2* (36, 60). Of note, in the previous study with AdV-mediated TBX18 overexpression, the amplitude of *I*_K1_ decrease is much higher than that of *I*_f_ increase, suggesting the major cause of the reported Ad-TBX18-induced pacing activity might come from *I*_K1_ suppression (11). The EAD-like oscillations are likely the result of a decrease in *I*_K_, resulting in long APs with repolarization disorders.

Previous studies have shown TBX18-mediated biological pacing in rats, guinea pigs and pigs when TBX18 was injected at the apex (20), AV junction (15) or His-bundle (16), where the cardiac conduction system is substantially present. Cells within the cardiac conduction system express *Hcn4*, have potential automaticity and display a spindle-shaped morphology distinct from ventricular myocytes (61). Expressing TBX18 in these cells may lead to an increased beating rate due to the intrinsic pacemaker properties of Purkinje cells, and will be responsive to beta-adrenergic stimulation due to the presence of Hcn4. This could potentially explain the discrepancies observed across different studies.

In the presence of beta-adrenergic stimulation, we observed robust biological pacemaker activity in animals injected with AAV-cTnT-Hcn2. To the best of our knowledge, this is the first comprehensive *in vivo* demonstration of an AAV-based biological pacemaker. Functional biological pacemaker engineering via overexpression of ion channels and constitutively active mutants has been extensively studied using AdV vectors (6–9, 36). However, the induction of long-term function with such interventions critically relies on sustained transgene overexpression, which now seems to be feasible with AAV-mediated gene transfer.

In conclusion, ectopic TBX18 expression in cardiomyocytes leads to cytotoxicity and fibrosis, while TBX18 expression at non-cytotoxic levels suppresses several working myocardial genes. Nevertheless, it does not induce a pacemaker gene program, nor does it generate or enhance a biological pacemaker. On the other hand, biological pacemaker outcomes of AAV-cTnT-Hcn2 were robust, provided that isoproterenol was present. This outcome represents a highly encouraging finding, supporting further exploration and testing of Hcn2-based dual transgene strategies (6–8).

## Methods

### Sex as a biological variable

Our study exclusively examined female mice and rats. It is unknown whether the findings are relevant for male mice and rats.

### Vector preparation

#### AAV production

The transfer plasmids used for AAV production in this study were all derived from pAAV-CMV-GFP (Penn Vector Core, University of Pennsylvania) which contains AAV serotype 2 inverted terminal repeats (ITRs). Coding sequences of human *TBX18* with C-terminal FLAG tag and murine *Hcn2* with N-terminal FLAG tag were synthesized and cloned into AAV transfer plasmids. Primers containing FLAG sequence were sued to amplify GFP, which was then cloned into to the backbone. Oligos with sequence GGCCGGCCGCCATGGCCAGTTTCTACCGGT and GGCCACCGGTAGAAACTGGCCATGGCGGCC were annealed and inserted in front of transgene coding sequence to introduce an upstream open reading frame. AAV6 vectors were produced by double transfection of HEK293T. Low passage HEK293T cells were plated in thirty 145 mm dishes at the density of 1.5 × 10^7^ cells per dish in DMEM-GlutaMax (ThermoFisher Scientific) containing 10% fetal bovine serum (FBS) (Sigma) and 1% penicillin-streptomycin (P/S) (ThermoFisher Scientific). For AAV6 vectors, cells were transfected next day with 33 µg pDP6 and 16 µg AAV transfer plasmids per dish using linear polyethylenimine (PEI) (Polysciences Inc). Medium was replaced during transfection to DMEM containing 1% P/S. Three days after transfection, cells were collected by centrifugation and the medium was concentrated by tangential flow filtration using the ÄKTA flux s system (GE Healthcare) to a final volume of 20 mL. Cells and medium were then combined, frozen and thawed twice followed by DNAseI, Rnase A, and Benzonase treatment. AAV vectors were purified by iodixanol density-gradient ultracentrifugation overnight. The AAV-containing fraction was then collected and concentrated to 1 mL by buffer exchange to PBS containing 0.001% Pluronic F68 using Amicon Ultra-15 100 kDa centrifugal filter units (Millipore). Concentrated AAV vectors were aliquoted and stored at -80°C until use.

#### AdV production

AdV-TBX18-GFP vector used in *in vitro* experiment was purchased from abmgood (Richmond, Canada). AdV vectors used in animal experiments were produced in Revvity Gene Delivery GmbH (Graefelfing, Germany). The coding sequence of GFP and N-terminal FLAG tagged human TBX18 was cloned into a pO6 shuttle plasmid and transferred via recombination in a BAC vector containing the genome of a replication deficient Ad5-based vector deleted in E1/E3 genes as previously described by Ruszics et al (62). Recombinant viral DNA was released from the purified BAC-DNA by restriction digest with Pac I and transfected into HEK293 cells. Viral particles were released via freeze/thaw treatment and amplified on HEK293. Final release of viral particles was performed via Na-Deoxycholate treatment followed by CsCl ultracentrifugation and buffer exchange on PD10 columns. Infectious units (IU) were determined via immunhistochemical detection of the adenoviral hexon protein in HEK293 cells infected with serial dilutions using anti hexon antibody (Santa Cruz).

### NRVM isolation and transduction

NRVMs were isolated from 1-to 2-day-old rats as described previously (63). NRVMs were cultured in Tung medium, which is M199 containing (in mmol/L) NaCl 137, KCl 5.4, CaCl_2_ 1.3, MgSO_4_ 0.8, NaHCO_3_ 4.2, KH_2_PO_4_ 0.5,Na_2_HPO_4_ 0.3, and supplemented with 20 units/100 mL penicillin, 20 µg/100 mL streptomycin, 2 μg/100 mL vitamin B12, and 2% or 10% fetal bovine serum (FBS). Tung medium containing 10% FBS (10% Tung) was only sued on the first day. Cells were cultured in monolayers or in low density single cell preparations on fibronectin-coated plates at 37°C in 5% CO_2_. AAV vector transduction was performed at multiplicity of infection (MOI) = 10,000 in 2% Tung medium. AdV vector transduction was performed at MOI = 1 in culture media. Medium was changed to 2% Tung medium the day after transduction, and afterwards was refreshed every 2 days till analyses.

### Western blot

Cells were washed twice with cold Dulbecco’s PBS and incubated with RIPA buffer supplemented with cOMPLETE protease inhibitor (Roche) on ice for 2 min. Cell lysates were gather using cell scraper, collected and transferred to a microcentrifuge tube. Cell debris was removed by centrifugation and the supernatant was used for further analysis. Protein concentration was determined using Pierce BCA Protein Assay Kit (Thermo Scientific) according to manufacturer’s manual. Two and twenty ug of the cell lysates were loaded onto each well in the HEK and NRVM experiments, respectively. Cell lysates were separated by 4–20% precast polyacrylamide gel (Bio-Rad) and transferred to PVDF membranes (Bio-Rad). After blocking with 5% skim milk in Tris-buffered saline with Tween 20 (5% skim milk/TBST) for one hour at room temperature, membranes were incubated with Rabbit-anti-FLAG (Sigma, F7425, 1:500) and Mouse-anti-B-actin antibodies (Sigma, A5441, 1:5000) in 5% skim milk/TBST overnight at 4°C. Next day, membranes were washed twice with TBST and incubated with Donkey-anti-Rabbit (Sigma, NA9340, 1:5000) and Sheep-anti-Mouse (Sigma, NA9310, 1:5000) antibodies for 2 hours at room temperature. Membranes were washed twice with TBST and once with water, followed by chemiluminescence detection using Amsersham ECL detection reagents (Cytiva).

### *In vivo* gene delivery in mice

Six-to eight-week old female FVB mice were used for experiments. Mice were injected subcutaneously with buprenorphine (0.075 mg/kg) and carprofen (0.05 mg/kg) for analgesia, at least 30 min prior to surgery. Anesthesia was induced with 4% isoflurane in 1 L/min O_2_. Mice were shaved, intubated and placed on a heating mat to maintain body temperature. Subsequently, an analgesic mixture consisting of lidocaine (2 mg/kg) and bupivacaine (3 mg/kg) was applied subcutaneously at the site of the incision. Anesthesia was maintained using ventilation with 2% isoflurane in 1 L/min O_2_. Left thoracotomy was performed to expose the apex of the heart at the fourth intercostal space. To inject the vector into the apex, a 10 µL Hamilton syringe fitted with a 31-G needle (13 mm, point style 4) was inserted from the anterior LV towards the apex. Five µL of the viral vector solution was slowly administered at each injection site to administer a total volume of 20 µL viral vectors containing 1 × 10^11^ viral genomes. Mice were randomly selected to receive control or experimental vectors. The thoracotomy and skin were closed with a C-1 12 mm cutting needle with a 6-0 silicone coated braided silk wire (Sofsilk, Covidien). Post-surgery analgesia consisted of 4 days of *ad libitum* carprofen (Rimadyl Cattle, 0.06 mg/mL) in drinking water and wet food.

### *In vivo* gene delivery in rats

Six-month old female Sprague Dawley rats were used for experiments. Anesthesia was induced with 5% isoflurane in 2 L/min O_2_. Rats were shaved, intubated and placed on a heating mat to maintain body temperature. Meloxicam (5 mg/kg) and Buprenorphine SR (1 mg/kg) were delivered subcutaneously for analgesia. Anesthesia was maintained using ventilation with 2% isoflurane in 2 L/min O_2_. Partial right thoracotomy was performed to create an ambulatory complete AV-block model, as previously reported.(64) Briefly, the right atrial appendage was ligated and retracted to expose the AV node region with its characteristic fat pad. Monopolar electrosurgical current was delivered subepicardially to the AV nodal region via a sharp needle to ablate the region. Monopolar electrosurgical current was delivered subepicardially to the AV nodal region via a sharp needle to ablate the region at five watts for 60 sec. If AV block was not generated, the current delivery was repeated under the same condition. ECG was taken one week after to confirm stable complete AV-block. Upon validating stable complete AV-block, rats were randomly selected to receive control or experimental vectors. A partial left thoracotomy was performed to expose the apex of the heart at the fourth intercostal space and inject 1 × 10^9^ viral genomes AdV vectors or 2.5 × 10^11^ viral genomes AAV vectors, suspended in 50 µL. Control groups were injected with equivalent volume of saline. For rats injected with AAV vectors and its saline control, 3 mg/kg rapamycin was intraperitoneally injected every other day to control the immune response and improve the transduction efficiency. ECGs were recorded for 60 minutes one week after vector injection. Isoproterenol was injected intraperitoneally at a dose of 1 mg/kg, 15 min after the start of recording.

### RNA isolation, cDNA synthesis, Reverse-transcription PCR and RNA sequencing

Cells were collected in lysis buffer RP1 (Macherey-Nagel). Tissues were stored immediately in RNAlater, and at a later timepoint homogenized using tissue homogenizer. RNA was isolated using Nucleospin RNA kit (Macherey-Nagel) according to the manufacturer’s protocol. For RT-PCR, complementary DNA library was transcribed from 500-1000 ng total RNA with oligo-dT primers (125 µmol/L) and the Superscript II system (Invitrogen). RT-qPCR was performed using the LightCycler 480 Real-Time PCR system (Roche). For comparison across different treatment groups, relative expression of the transgene mRNA was normalized to the geomean of reference genes *Hprt1* and *Rpl4*. Fold change was calculated over the mean of the control group. For *Hcn4* expression over different time points, relative expression of the transgene mRNA was normalized to the reference gene *Hprt1* and fold change was calculated over day 0. For RNA sequencing, cDNA was prepared with KAPA mRNA Hyperprep Kit (Roche, KK8580). Total RNA (1 µg) was purified and sheared into small fragments followed by cDNA synthesis and ligation of the adapter. The quality and size distribution of the cDNA library templates were validated with Agilent DNA 1000 on 2100 Bioanalyzer (Agilent Technologies). cDNA samples were pooled per lane of paired-end 150 bp sequencing on a NovaSeq X Plus instrument (llumina).

### Differential expression analysis

Reads were trimmed using Trim Galore! (https://https://github.com/FelixKrueger/TrimGalore.com/fenderglass/Flye) and mapped to rn6 build of the rat transcriptome or mm10 build of the mouse transcriptome using STAR (65). Differential expression analysis was performed using DESeq2 (66). Unsupervised hierarchical clustering was performed on genes differentially expressed (between X, Y and Y) using the R package pheatmap, version 1.0.12. (http://cran.r-project.org/web/packages/pheatmap/index.html). Gene Ontology (GO) terms and Kegg pathways were found using DAVID (67). For the comparison to Goodyer et al., the fold change of insignificantly differentially expressed genes were set to 1 where adjusted p-value < 0.01 was considered significant.

### Histological staining

#### Immunofluorescence staining

Hearts were harvested and fixed in 4% PFA overnight, embedded in paraffin and sectioned at 7 µm thickness. Sections were deparaffinized and dehydrated by a series of ascending ethanol concentrations. For antigen retrieval, sections were boiled in unmasking solution (H3300, Vector). Sections were blocked in 4% bovine serum albumin (BSA) and incubated in 2% BSA with nuclear staining and primary antibodies. The primary antibodies used were: chicken anti-GFP (1:500, Aves Labs, GFP-1020), rabbit anti-FLAG (1:200, Millipore, F7425), Goat anti-TBX18 (1:50, Santa Cruz, SC-17869) and DAPI (1 µg/mL, Sigma, D9542). Fluorescence images were acquired using a Leica DM6000 fluorescence microscope or a Leica TCS SP8 confocal microscope (Leica Microsystems).

#### Picrosirius red staining

Hearts were harvested and fixed in 4% PFA overnight, embedded in paraffin and sectioned at 7 µm thickness. Sections were deparaffinized and dehydrated by a series of ascending ethanol concentrations and stained with picrosirius red. Images were acquired using a Leica DM5000 fluorescence microscope (Leica Microsystems). Image analysis was performed on 4 non-contiguous sections from the anterior of the heart ImageJ. Total area of the left ventricle and the area of fibrosis were measured from which the percentage of fibrosis was calculated.

### Patch clamp experiments

#### Cell preparation and data acquisition

Left ventricular cardiomyocytes of mice were isolated by an enzymatic dissociation procedure (68). Cells were stored at room temperature for at least 45 min in a modified Tyrode’s solution containing (in mmol/L): NaCl 140, KCl 5.4, CaCl_2_ 1.8, MgCl_2_ 1.0, glucose 5.5, HEPES 5.0; pH 7.4 (NaOH) before use. Cells were put into a recording chamber on the stage of an inverted microscope (Nikon Diaphot, Melville, New York, NY, USA), and single GFP-positive quiescent mouse cardiomyocytes with smooth surfaces were selected for electrophysiological measurements. Action potentials (APs) and net membrane currents were recorded with the amphotericin-perforated patch-clamp technique, while the sodium current (*I*_Na_) was measured with the ruptured patch clamp, using an Axopatch 200B amplifier (Molecular Devices, Sunnyvale, CA, USA). Pipettes (resistance 2–3 MΩ) were pulled from borosilicate glass capillaries (TW100F-3; World Precision Instruments) and filled with solutions as indicated below. Voltage control, data acquisition, and analysis were realized with custom software (69). Signals were low-pass-filtered with a cutoff of 5 kHz and digitized at 40 kHz (APs), 5 kHz (net currents), and 20 kHz (*I*_Na_). Potentials were corrected for the calculated liquid junction potential.(70) Cell membrane capacitance (C_m_) was estimated by dividing the time constant of the decay of the capacitive transient in response to 5 mV hyperpolarizing voltage clamp steps from –40 mV by the series resistance (69). For net currents and *I*_Na_ measurements, C_m_ and series resistance were compensated for by at least 60 and 80%, respectively.

#### AP measurements

APs were recorded at 36 ± 0.2°C using the modified Tyrode’s solution as bath solution. Pipette solution contained (in mmol/L): K-gluconate 125, KCl 20, NaCl 5.0, amphotericin-B 0.44, HEPES 10, and pH 7.2 (KOH). The single cells from dtTBX18-GFP positive cells were spontaneously beating. We measured spontaneously APs as well as APs APs elicited at an overdrive stimulation of 4 Hz by 3-ms, ≈20–40% suprathreshold current pulses, through the patch pipette. Averages AP parameters of stimulated cells were taken from 10 consecutive APs and we analyzed the maximal diastolic potential (MDP), maximum AP upstroke velocity (V_max_), AP amplitude (APA), AP plateau potential (APplat), and APD at 20, 50, and 90% repolarization (APD_20_, APD_50_, and APD_90_, respectively). Parameters from 10 consecutive APs were averaged.

#### Net membrane current measurements

APs measurements were followed by voltage clamp experiments to determine which changes in currents underlay the AP differences between WT and TBX18 overexpressing cells. To make a direct comparison between MDP and background K^+^ currents within one cell, and to ensure that AP shapes and the remaining viable cardiomyocytes in the recording chamber stay undistorted for biophysical analysis, these measurements were performed without specific channel blockers or modified solutions. Membrane currents were examined by 500 ms voltage-clamp steps every 2 s to membrane potentials ranging from -100 to +50 mV from a holding potential of -40 mV. The inward rectifier K^+^ current (I_K1_) and delayed rectifier K^+^ current (I_K_) were defined as the quasi steady-state current at the end of the voltage-clamp steps at potentials negative or positive to -30 mV, respectively. The transient outward K^+^ currents (I_to’s_) was defined as the difference between the peak inward current and the current amplitude at the end of the 500 ms depolarizing voltage clamp step. All currents were normalized for C_m_. Presence of the hyperpolarization-activated, funny current (I_f_) was tested with 2000 ms voltage-clamp steps every 5 s to membrane potentials ranging from -120 to -40 mV from a holding potential of -40 mV. Differences between the current at the begin and the current at the end of the hypolarizing voltage clamp step was defined as I_f_.

#### I_Na_ measurements

*I*_Na_ *was* characterized at room temperature using a bath solution containing (in mmol/L): NaCl 7, CsCl 133, CaCl_2_ 1.8, MgCl_2_ 1.2, glucose 11.0, HEPES 5.0, and pH 7.4 (CsOH). Nifedipine (5 µmol/L) was added to block the L-type calcium current. Pipettes were filled with solution containing (in mmol/L): NaCl 3, CsCl 133, MgCl_2_ 2.0, Na_2_ATP 2.0, TEA-Cl 2.0, EGTA 10.0, HEPES 5.0, and pH 7.2 (CsOH). *I*_Na_ amplitudes and voltage dependency of *I*_Na_ activation were measured by 50 ms depolarizing voltage clamp steps from a holding potential of -120 mV. A two-step protocol consisting of series of 500 ms pulses between -140 and 40 mV from a holding potential of -120 mV, followed by a second 50 ms step to -20 mV, was used to establish the voltage dependency of *I*_Na_ inactivation. Cycle lengths of the protocols were 5 s. *I*_Na_ was defined as the difference between peak current and steady-state current and *I*_Na_ density was calculated by dividing current amplitudes by C_m_. Voltage dependence of activation and inactivation were determined by fitting a Boltzmann function (y=[1+exp{(V-V1/2)/k}]^-1^) to the individual plots, where V_1/2_ is the voltage of half-maximal (in)activation and k the slope factor (in mV).

### Statistical analysis

GraphPad Prism version 10 software (GraphPad Software Inc.) was used for statistical analysis. Numerical data are presented as mean ± standard error of the mean (SEM) of at least three biological replicates. Two independent groups were compared by Mann-Whitney test and multiple independent groups were compared by analysis of variance (ANOVA) followed by the *post-hoc* Holm-Šídák test or Fisher’s LSD test for multiple comparisons. Categorical data were presented as absolute values and percentages. Categorical data were compared by Fisher’s exact test. All tested were performed two-sided and statistical significance was considered for p < 0.05.

### Study approval

Amsterdam UMC (the Netherlands): All mice were treated in accordance with the EC Directive 2010/63/EU of the European Parliament and of the Council on the protection of animals used for scientific purposes and using protocols approved and monitored by the Animal Welfare Body of Amsterdam UMC.

Naason Science Inc. (Republic of Korea): Rat study was carried out according to Korea Food and Drug Administration guidelines for the care and use of laboratory animals and approved by the Institutional Animal Care and Use Committee.

## Supporting information

Supplemental materail

## Data availability

The datasets generated or analyzed during the studies reported herein are available in the Supporting Data Values file provided in a single Excel sheet, and repeats and supporting data sets can be obtained from the corresponding author on reasonable request.

## Author contribution

JW designed and conducted experiments, analyzed data, drafted the manuscript, and oversaw the project. MRR, MK, ARB, RNV, RS, HYL and KvD contributed to the conduct of experiments. KHP and LCP oversaw the conduct of rat experiments. SS, CT, EE and AE contributed to the vector production. OFK, KN and HLT designed experiments and revised the manuscript. AOV designed, conducted experiments, and analyzed data. VM and GJJB designed experiments, drafted and revised the manuscript, and oversaw the project.

## Acknowledgement

This work is supported by funding from The European Research Council starting grant 714866 and associated proof-of-concept grant 899422 (to G.J.J.B.), Health Holland LentiPace II (to G.J.J.B. and H.L.T.), Horizon 2020 Eurostars (E114245 and E115484; to G.J.J.B. and V.M.C.); Dutch Research Council Open Technology Program 18485 (to H.L.T. and G.J.J.B.); The European Innovation Council (EIC; TRANSITION - Project 101099608 – TRACTION; to V.M.C. and G.J.J.B.); and The Dutch Research Council OCENW.GROOT.2019.029 (to V.M.C.). We would like to thank Otto J. Mulleners for his help in RNA sequencing analysis.

