## Supplemental materail for "Adeno-associated virus (AAV)-TBX18 does not generate biological pacemaker activity, unlike AAV-Hcn2"

### Supplemental material

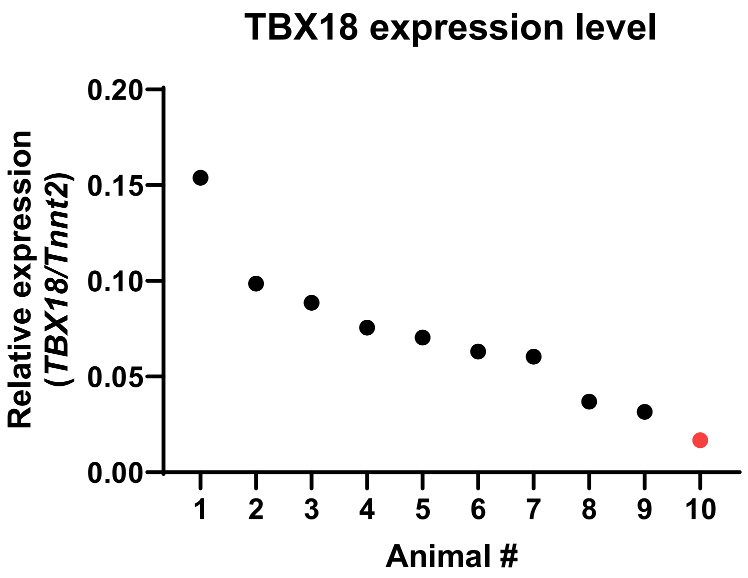


**Supplemental Figure S1 TBX18 expression level in mouse hearts injected with AAV-TBX18  4 weeks post injection.** Red dot indicates the single TBX18 heart in which no fibrosis was detected.

A

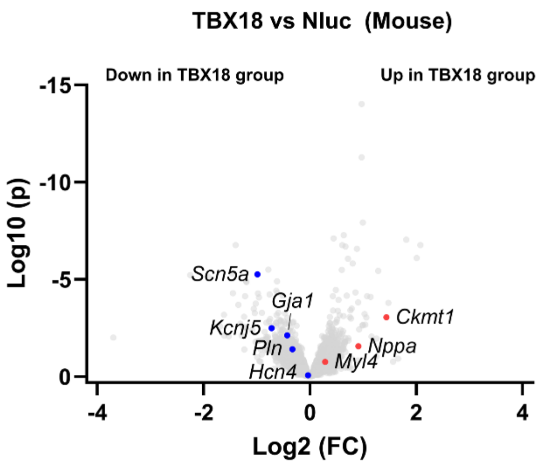


B

| Enriched GO term in down-regulated genes | p-adjust |
|---|---|
| anatomical structure development (GO:0048856) | 1.62E-10 |
| developmental process (GO:0032502) | 1.47E-09 |
| cellular developmental process (GO:0048869) | 5.83E-08 |
| cell differentiation (GO:0030154) | 7.69E-08 |
| regulation of multicellular organismal process (GO:0051239) | 3.39E-07 |
| cell development (GO:0048468) | 4.15E-07 |
| multicellular organismal process (GO:0032501) | 4.86E-07 |
| monoatomic cation transmembrane transport (GO:0098655) | 5.50E-07 |
| inorganic ion transmembrane transport (GO:0098660) | 1.33E-06 |
| monoatomic ion transmembrane transport (GO:0034220) | 1.52E-06 |

| Enriched GO term in up-regulated genes | p-adjust |
|---|---|
| negative regulation of nitrogen compound metabolic process (GO:0051172) | 1.35E-02 |
| circulatory system process (GO:0003013) | 1.44E-02 |
| anatomical structure development (GO:0048856) | 1.47E-02 |
| developmental process (GO:0032502) | 1.47E-02 |
| negative regulation of cell communication (GO:0010648) | 1.54E-02 |
| sensory perception of smell (GO:0007608) | 1.60E-02 |
| cellular developmental process (GO:0048869) | 1.65E-02 |
| regulation of release of cytochrome c from mitochondria (GO:0090199) | 1.68E-02 |
| cellular process (GO:0009987) | 1.69E-02 |
| negative regulation of signaling (GO:0023057) | 1.69E-02 |

**Supplemental Figure S2 TBX18 expression in cardiomyocytes suppressed chamber myocardial genes.**

(**A**) Volcano plot presenting gene changes between mouse hearts injected with AAV-cTnT-uORF-Nluc-GFP and AAV-cTnT-uORF-TBX18-GFP. (**B**) Gene ontology analysis on the most significantly up- and down-regulated genes in mouse hearts injected with AAV-cTnT-uORF-TBX18-GFP.

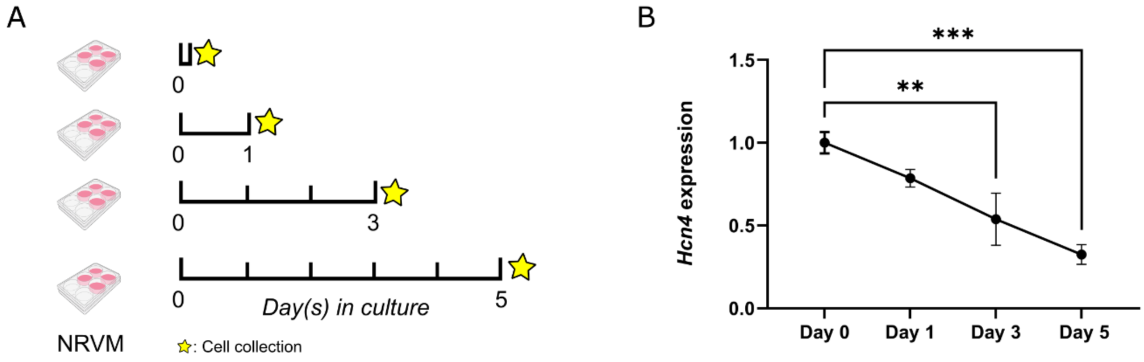


**Supplemental Figure S3 *Hcn4* expression decreases over time in NRVMs.**

(**A**) Experimental design. (**B**) Expression level of *Hcn4* in NRVMS at 0, 1, 3 and 5 days in culture. Data are shown as mean ± SEM. Data were compared using one-way ANOVA with *post-hoc* Fisher's LSD test. $^{**}p < 0.01$; $^{***}p < 0.001$.

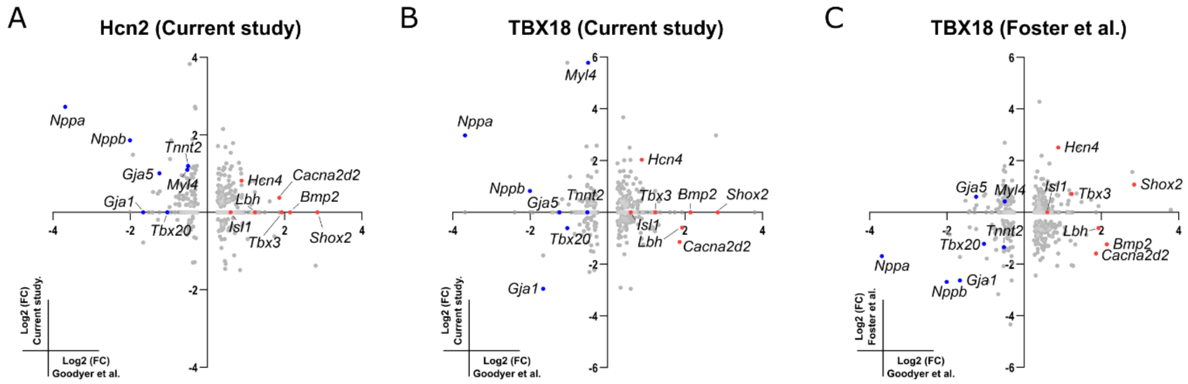


**Supplemental Figure S4 Scatter plots showing the fold change of SAN marker genes selected from Goodyer et al. in various RNA-seq datasets.**

(**A**) NRVM transduced with Hcn2 from current study. (**B**) NRVM transduced with TBX18 from the current study. (**C**) NRVM transduced with TBX18 from Foster et al.
